# Understanding and evaluating ambiguity in single-cell and single-nucleus RNA-sequencing

**DOI:** 10.1101/2023.01.04.522742

**Authors:** Dongze He, Charlotte Soneson, Rob Patro

## Abstract

Recently, a new modification has been proposed by Hjörleifsson and Sullivan *et al*. to the model used to classify the splicing status of reads (as spliced (mature), unspliced (nascent), or ambiguous) in single-cell and single-nucleus RNA-seq data. Here, we evaluate both the theoretical basis and practical implementation of the proposed method. The proposed method is highly-conservative, and therefore, unlikely to mischaracterize reads as spliced (mature) or unspliced (nascent) when they are not. However, we find that it leaves a large fraction of reads classified as ambiguous, and, in practice, allocates these ambiguous reads in an all-or-nothing manner, and differently between single-cell and single-nucleus RNA-seq data. Further, as implemented in practice, the ambiguous classification is implicit and based on the index against which the reads are mapped, which leads to several drawbacks compared to methods that consider both spliced (mature) and unspliced (nascent) mapping targets simultaneously — for example, the ability to use confidently assigned reads to rescue ambiguous reads based on shared UMIs and gene targets. Nonetheless, we show that these conservative assignment rules can be obtained directly in existing approaches simply by altering the set of targets that are indexed. To this end, we introduce the *spliceu* reference and show that its use with `alevin-fry` recapitulates the more conservative proposed classification.

We also observe that, on experimental data, and under the proposed allocation rules for ambiguous UMIs, the difference between the proposed classification scheme and existing conventions appears much smaller than previously reported. We demonstrate the use of the new `piscem` index for mapping simultaneously against spliced (mature) and unspliced (nascent) targets, allowing classification against the full nascent and mature transcriptome in human or mouse in <3GB of memory. Finally, we discuss the potential of incorporating probabilistic evidence into the inference of splicing status, and suggest that it may provide benefits beyond what can be obtained from discrete classification of UMIs as splicing-ambiguous.

## 1. Introduction

The evaluation of the splicing status of molecules, as spliced (mature) or unspliced (nascent), is an important task in the analysis of single-cell and single-nucleus RNA-seq data. Popular droplet-based technologies, like the 10x Genomics Chromium platform, often sequence a mixture of spliced (mature) and unspliced (nascent) RNA sequences (1), and recent work has suggested that there may be benefits to accounting for UMIs arising from both spliced (mature) and unspliced (nascent) molecules (1–3) in subsequent analysis. Moreover, it may often be important to separate the counts of molecules of different splicing status, allocating the associated UMIs as having a spliced or unspliced (or sometimes ambiguous) origin. For example, analyses based on splicing status-aware counts have been used to study (and simulate) cell differentiation and development processes (4–9) and have proven useful in other important research endeavors, such as disease state prediction (10). Therefore, it is important to understand, analyze, and improve the methods by which the splicing status of UMIs is inferred.

Recently, Hjörleifsson and Sullivan *et al*. (11) introduced a new strategy for assigning reads (and subsequently UMIs) to splicing states and genes in single-cell and single-nucleus RNA-seq data. In this new scheme, they propose to classify reads as unspliced (nascent) (N), spliced (mature) (M), or ambiguous (N|M) in origin — an existing classification scheme (4, 5), for which Hjörleifsson and Sullivan et *al*. (11) adopt different definitions. Under the proposed classification scheme, the groups are defined as follows: (i) unspliced (nascent) UMIs are those that arise from reads contained within intronic sequences or spanning intron-exon boundaries, (ii) spliced (mature) UMIs are those that arise from reads spanning exon-exon junctions, and (iii) ambiguous UMIs are those that are neither nascent nor mature (typically, those that arise from reads contained entirely within an exon). The proposed scheme is conservative compared to existing classification approaches, because reads deriving entirely from exons are recognized as ambiguous in origin, whereas they are typically classified as mature in existing approaches (4, 5, 12, 13).

While the conservative motivation of this classification scheme makes sense conceptually, its current practical utility is questionable. Specifically, in a typical experiment (either single-cell or single-nucleus RNA-seq), a large fraction of all reads will be classified as ambiguous (**Table 1)**. On average, existing assignment rules allocate ~ 10% of the UMIs as ambiguous for the datasets evaluated in **Table 1,** so the proposed scheme attributes about five times as many UMIs to the ambiguous category compared to existing assignment rules. In fact, this reality is implicitly acknowledged in the manuscript itself. The authors perform a set of experiments: (i) evaluating gene abundance estimation accuracy on simulated single-cell RNA-seq data, (ii) performing a “classification” experiment on simulated reads from molecules of different splicing statuses, and (iii) comparing a variant of the proposed approach to existing pipelines in the processing of single-nucleus RNA-seq data. Yet, in each of these experiments, the UMIs falling into the ambiguous class are allocated *differently* and precisely in the manner that will most benefit the proposed method in the assessment being carried out (i.e., maximizing the accuracy in the simulated experiments and highlighting the difference of the proposed approach and existing classification schemes in the processing of an experimental sample). As such, this approach raises several immediate concerns. Below, we highlight several conceptual concerns, suggest improvements or alternatives, and also report a likely evaluation error in the results of Hjörleifsson and Sullivan *et al*. (11) that led to an over-stated difference among methods.

**Table 1.**
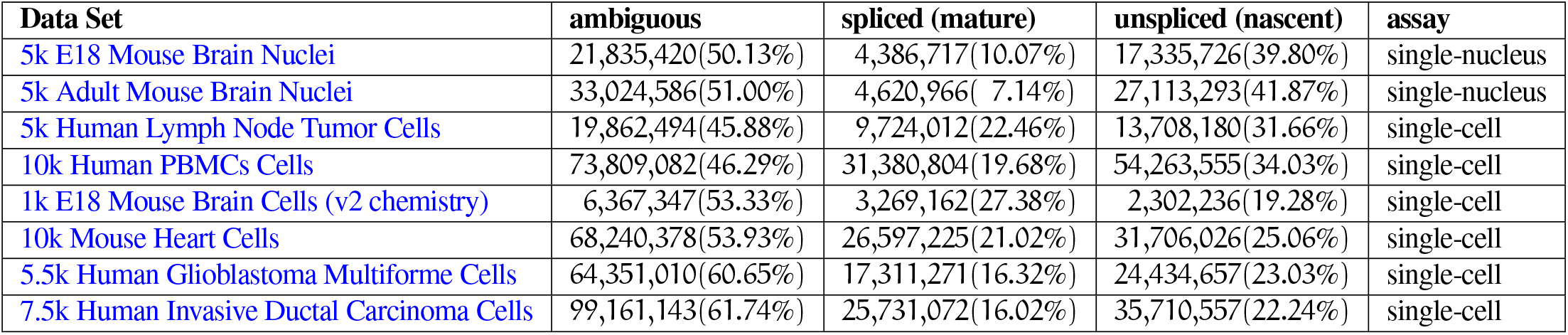
The number of spliced (mature), unspliced (nascent), and ambiguous UMIs, as classified under the model proposed by Hjörleifsson and Sullivan *et al*. (11), across 8 single-cell and single-nucleus data sets using the 10x Chromium platform. The percentages are calculated as 100 times the total number of UMIs in each category, divided by the total number of assigned UMIs.

### Inconsistent treatment and allocation of ambiguous UMIs

Hjörleifsson and Sullivan *et al*. critique existing approaches based on the observation that they attribute reads (and subsequently UMIs) of exonic origin as spliced (mature). It is noted, correctly, that, in the strictest sense, these reads are ambiguous since they are compatible with both spliced (mature) and unspliced (nascent) RNA. The authors propose, instead, that such UMIs should be considered as ambiguous. However, the authors do not consistently apply this definition in their manuscript. For example, in the classification experiment (Figure 2 of Hjörleifsson and Sullivan *et al*. (11)) these UMIs are actually classified as ambiguous, but in the simulated single-cell experiment, the ambiguous UMIs are all allocated to spliced (mature) molecules, and in the experimental single-nucleus sample, the ambiguous UMIs are all allocated to unspliced (nascent) molecules. Thus, the authors choose, in practice, to allocate these ambiguous UMIs differently, and in an all-or-nothing manner, in each context. These allocations are also made in the way that most benefits the assessed performance of their proposed method. Also, it is worth noting that the classification scheme of `alevin-fry` (13) is a result of the way the *splici* index is constructed, and is adopted to conform to prevailing convention (4, 5). In fact, by indexing gene-length unspliced (nascent) molecules instead of the collapsed introns (similar to what was explored in Soneson et al. (14), but with one unspliced molecule per gene instead), one immediately obtains a reference (a *spliceu* reference) that induces a classification scheme compatible with what is proposed by Hjörleifsson and Sullivan *et al*. (11) (see Section 2).

In reality, ambiguous UMIs are likely to come from a combination, *with unknown mixture proportions*, of both spliced (mature) and unspliced (nascent) RNAs. Further, these proportions can vary from sample to sample, cell type to cell type, across genes (3), and, of course between different assays. Simply classifying these UMIs as ambiguous, while a conservative approach, vastly reduces the amount of usable data under existing downstream processing pipelines, further exacerbating the challenges posed by the sparse sampling of the most prevalent single-cell sequencing technologies. Further, assigning them entirely to either a spliced or unspliced status, depending on the type of assay being processed, does little to ameliorate the problem beyond prevailing convention.

### Difficulty in reproducing certain results due to absent files and code

We observed that the “significantly different” (11) results of the proposed approach, highlighted when quantifying an experimental single-nucleus RNA-seq sample, cannot be easily reproduced. The scripts provided by Hjörleifsson and Sullivan *et al*. reference local file resources without providing instructions to generate these files and without making these files available. When using `kallisto(D-list)|bustools` to quantify the data against a nascent transcriptome constructed in a way consistent with the description provided by Hjörleifsson and Sullivan *et al*., under the proposed assignment rules, the quantification results are highly concordant with the existing tools (i.e. `STARsolo` and `alevin-fry`). Also, while the results from that manuscript cannot be directly reproduced without the referenced files, we observe that if the nascent transcriptome is extracted with the `generate_cDNA+introns.py` script provided by the authors, one obtains “significantly different” quantification results (broadly similar to those demonstrated by Hjörleifsson and Sullivan *et al*.) from both existing tools and from `kallisto(D-list)|bustools` constructed with the nascent transcriptome extracted according to the paper’s description. This difference appears to be due, almost entirely, to the fact that the `generate_cDNA+introns.py` script does not reverse complement nascent transcripts arising from genes on the negative strand prior to indexing. All analyses and references to the scripts of Hjörleifsson and Sullivan *et al*. (11) in this manuscript are with respect to commit cfd6958be6ab65fc340bf1df5f1b0e77f19ff967, which was the latest available when this manuscript was submitted.

### Discarding – rather than resolving and quantifying – fragments that arise from the unexpected (or less dominant) splicing status may result in misallocation of UMIs that could have been properly recovered

Surprisingly, the approach proposed by Hjörleifsson and Sullivan *et al*. does not actually categorize the splicing status of each UMI. Rather, the authors construct entirely separate indices (and associated D-lists) for the spliced (mature) and unspliced (nascent) references. The ambiguous category is then defined implicitly as those reads that map under both indices. As such, to fully categorize reads as definitively spliced (mature), definitively unspliced (nascent), or ambiguous, one must construct two separate indices, map the reads twice, and must then compare the output to determine the reads that should be treated as ambiguous. However, the authors do not provide a tool or pipeline for this final step. Likewise, no mechanism is proposed for how to deal with mapped fragments that may be ambiguous, but which share a UMI (and appear in the same cell barcode) with mapped fragments that are not ambiguous in origin. Therefore, in practice, by restricting the information available to the index and to the subsequent mapping and resolution algorithms, this approach may fail to properly allocate UMIs that could be properly allocated via mapping under a more comprehensive index.

This is important because spliced (mature) or unspliced (nascent) reads confidently sharing a UMI with ambiguous reads can ideally be used as evidence to avoid misallocating this UMI. See e.g. **Fig. 1.** Here, an unspliced (nascent) index is constructed using the spliced (mature) transcriptome as the D-list (i.e., to determine the distinguishing flanking k-mers (DFKs)). There are 2 reads (black bars) in the same barcode sharing a UMI. The leftmost read maps entirely within an exon, and is therefore attributed to the nascent count (under the rules proposed by Hjörleifsson and Sullivan *et al*. (11) it will be ambiguous, but the algorithm does not distinguish between purely nascent and exonic reads). The right read, however, maps partially within the same exon but overlaps a DFK to the right. Because this read includes a DFK, it will be discarded and not considered further in the quantification pipeline. However, this read actually provides strong evidence that the shared UMI should be attributed to the spliced (mature) version of the gene being quantified, rather than the unspliced (nascent) version, as the DFK arises from a splicing junction on a spliced (mature) transcript of this gene.

**Fig. 1.**
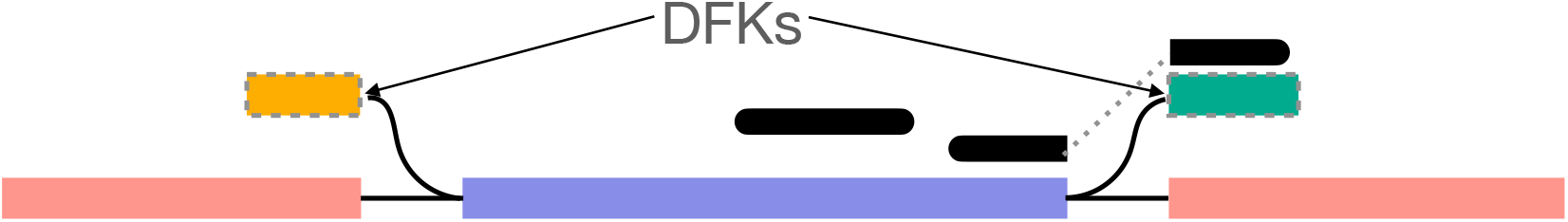
Discarding — rather than resolving and quantifying — the fragments that arise from the unexpected (or less dominant) splicing status in a sample may result in misallocation of UMIs that could have been properly recovered. Here, the sequenced fragments (black bars) share the same cell barcode and UMI. The leftmost fragment maps entirely within the (purple) exon, and is therefore quantified against the mapped gene in the underlying (unspliced (nascent)) index. The rightmost fragment maps across an exon-exon junction, and contains a distinguishing flanking k-mer (DFK); it is therefore discarded during mapping. Yet, because these fragments map to the same gene and share the same UMI, it is most likely that they both arise from the same (spliced) pre-PCR molecule, and that the associated UMI should be allocated toward the spliced version of this gene.

### A single index, paired with a splicing aware UMI resolution algorithm, can permit the efficient and explicit assignment of UMI splicing status

In addition to potentially improving UMI assignment, since the counts of UMIs arising from molecules of all splicing statuses may still be meaningful, for example, in RNA velocity (4, 5) and disease state prediction (10), it may be useful to have them mapped and quantified for subsequent analysis, regardless of the assay being processed. To obtain these counts using the `kallisto(D-list)|bustools` approach, one currently needs to construct and quantify against two separate indices, which is less efficient than constructing and quantifying against a single index. However, a single mapping and quantification run of `alevin-fry` (against either the *splici* or *spliceu* reference, depending on the ambiguity convention one wishes to adopt) or of STARsolo, is sufficient to quantify and classify all such reads. While the existing *splici* and *spliceu* indices of `alevin–fry` are already quite memory frugal — requiring ~ 8GB of memory for quantification against human and mouse-sized references using the existing `pufferfish` (15) index — here, we also introduce the new `piscem` (16) index and mapper, which further lowers the memory requirements for this type of analysis. When indexing the full spliced (mature) and unspliced (nascent) transcriptomes, the `piscem` (16) index is <2GB for human and mouse, and the read mapping using `piscem` itself can be carried out in ≤ 3GB for all of the experiments considered in this manuscript.

## 2. Results

To better understand the implications of the classification scheme proposed by Hjörleifsson and Sullivan *et al*. (11), we re-created several of the experiments described in that manuscript. Primarily, we were interested in further understanding the characteristics of the classification simulation and the nature of the observed differences in the experimental mouse single-nucleus RNA-seq dataset. In the classification experiment (underlying Figure 2 of Hjörleifsson and Sullivan *et al.)*, we sought to observe how an `alevin-fry` reference constructed to conform to their proposed classification rules (i.e., a *spliceu* reference) would perform under their assessment; these results are described in Section 2.1. In the mouse single-nucleus RNA-seq dataset analysis (underlying Figure 4 of Hjörleifsson and Sullivan *et al*.), we sought to understand the fraction of UMIs that would be classified as ambiguous under the proposed model, and how such a small fraction of confidently spliced (mature) UMIs in this data could result in such a seemingly large difference in quantification results; these results are described in Section 2.2. Additionally, in Section 2.3 we discuss the incorporation of probabilistic evidence into the inference of the splicing status of UMIs.

**Fig. 2.**
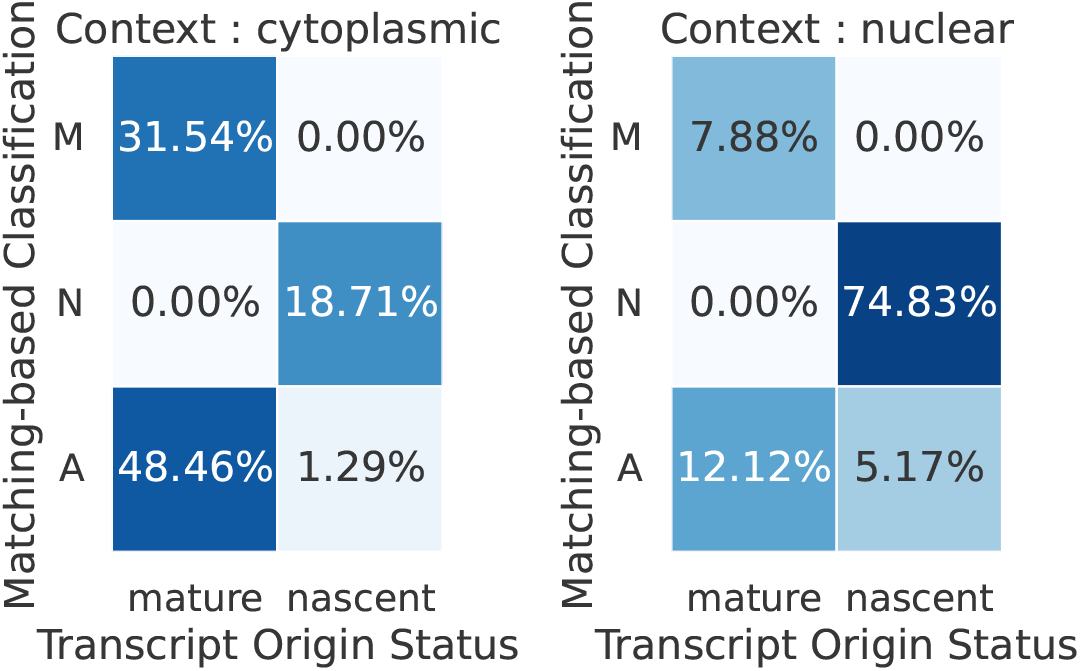
The frequency with which reads arising from a molecule of a specific splicing origin, either spliced (mature) or unspliced (nascent) (given on the x-axis), are assigned each status spliced (mature) (M), unspliced (nascent) (N), or ambiguous (A) under the proposed empirical assignment rules (given on the y-axis). These assignment frequencies are illustrated for simulations in both the cytoplasmic (left) and nuclear (right) contexts. A perfect assignment rule would classify all reads from spliced (mature) transcripts as mature, and would classify all reads from unspliced (nascent) transcripts as nascent, though sufficient information may not be available to make such a classification in practice, leading to the need for the ambiguous category. The difference between the cytoplasmic and nuclear contexts is the relative number of reads simulated from mature and nascent transcripts.

**Fig. 3.**
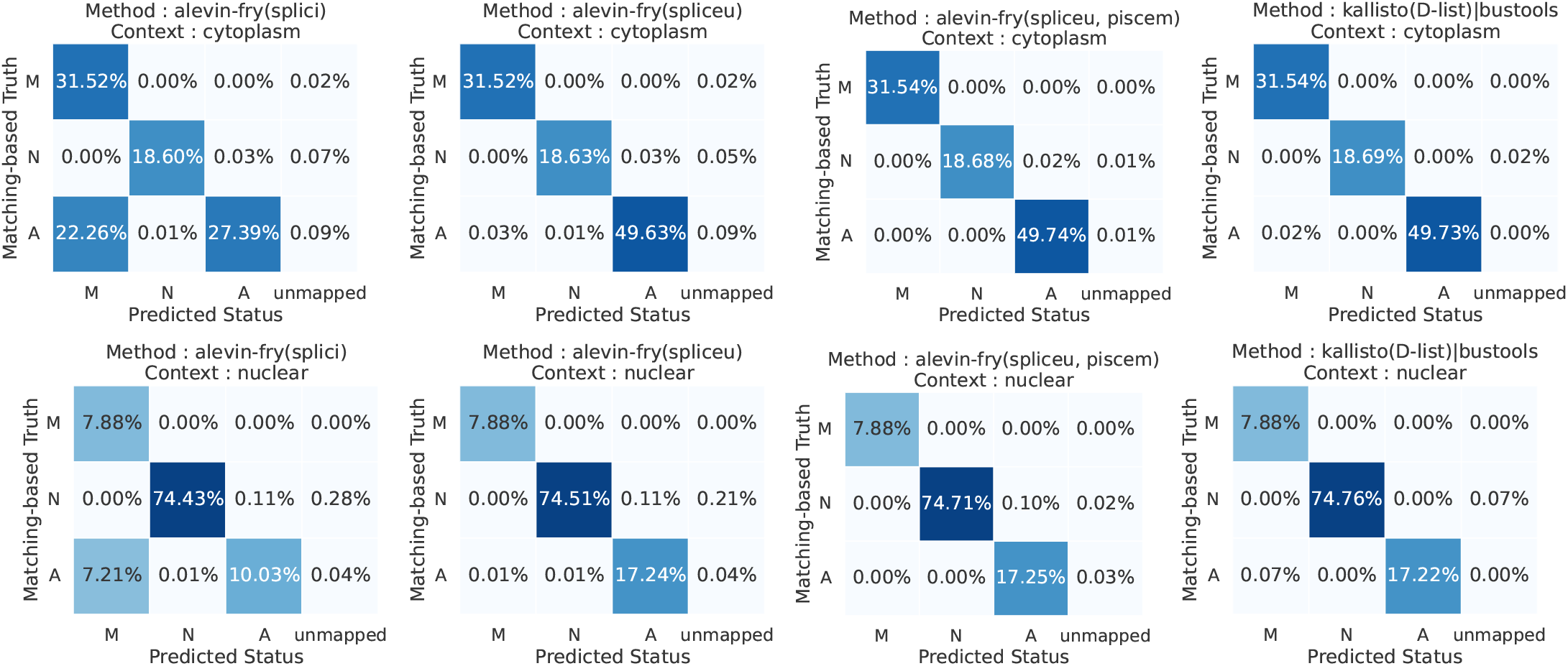
The assignment status for the simulated reads, as mapped by each method, in both the cytoplasmic context (top row) and nuclear context (bottom row). The x-axis of each heatmap enumerates the predicted status (under that tool) for the simulated reads, and the y-axis enumerates the “true” status (i.e., the matching-based status assessed under perfect matching of the reads to their compatible targets). The fourth column of each matrix shows the fraction of reads in each category that remained unmapped. The “Method” atop each matrix lists the method whose mappings are being evaluated. The agreement is assessed on the per-read level, and then classified and misclassified reads are aggregated to produce the statistics displayed in each matrix. A method with perfect *internal consistency* with the model being tested would allocate all reads along the diagonal, and no reads off-diagonal.

**Fig. 4.**
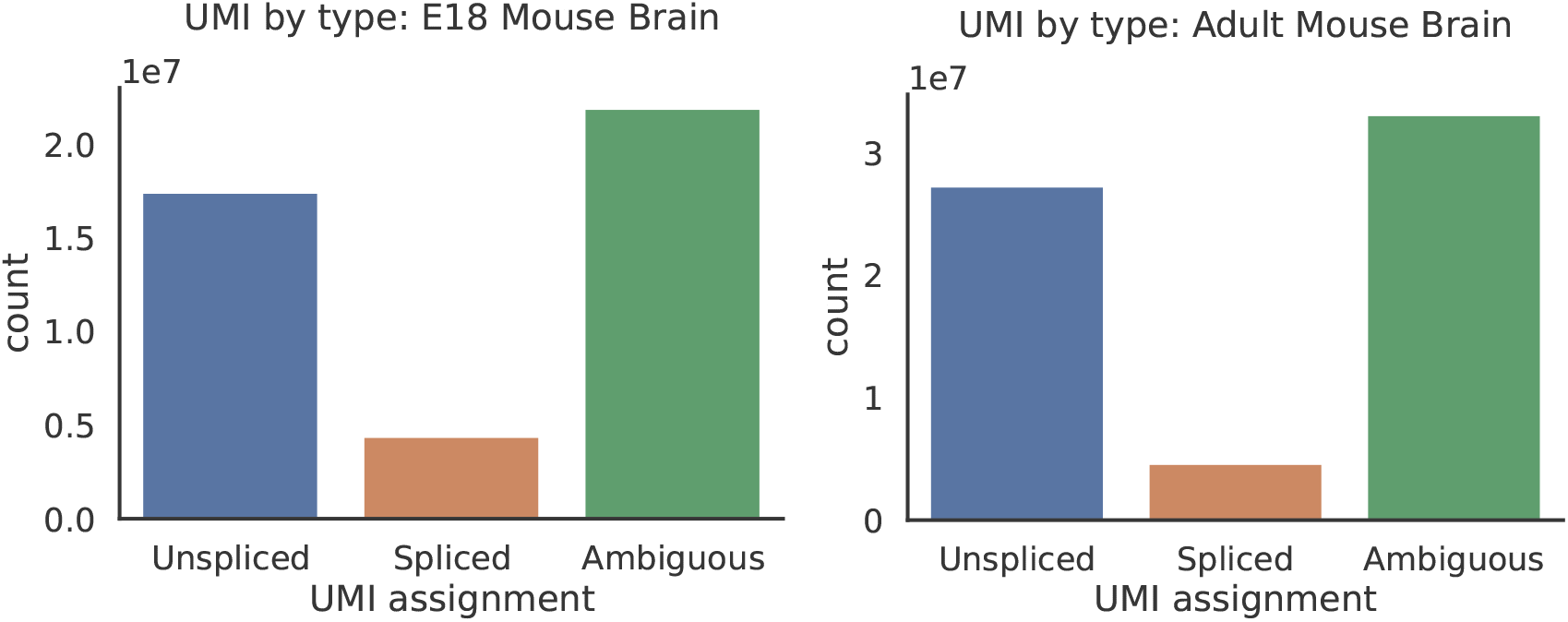
The count of UMIs confidently predicted to arise from unspliced, spliced molecules or to be ambiguous in splicing status, as characterized under the model proposed by Hjörleifsson and Sullivan *et al*. (11) in the E18 mouse brain nuclei dataset (left) and the adult mouse brain nuclei dataset (right).

### 2.1. Status classification of simulated data

We carried out a version of the classification experiment proposed by Hjörleifsson and Sullivan *et al*. (11), simulating reads from a mix of spliced (mature) and unspliced (nascent) transcripts from both cytoplasmic and nuclear contexts, and evaluated the inferred status of these reads under the mappings produced by different tools and indices. Specifically, we considered `kallisto(D-list)|bustools`, `alevin-fry(splici)`, `alevin-fry(spliceu)`, and `alevin-fry(spliceu, piscem)`. The latter uses the exact same *spliceu* reference as `alevin-fry(spliceu)`, but `alevin-fry(spliceu, piscem)` uses `piscem` instead of `salmon` (17) to perform the mapping.

First, we analyzed how the classification rules adopted in this model accord with the true splicing origin of each read (**Fig. 2)**. That is, every read truly originates from either a spliced (mature) or unspliced (nascent) molecule, and the ambiguous status acts as a convenient classification for a read when a definitive splicing status cannot be inferred from the mapping information alone. However, the utility of this ambiguous classification depends on how the ambiguous reads are ultimately allocated, as well as the fraction of ambiguous reads that truly arise from either spliced (mature) or unspliced (nascent) molecules. In these simulations, we observe a substantial asymmetry in the rate at which these reads are classified as ambiguous. Specifically, in both contexts, a large fraction of truly spliced (mature) reads are classified as ambiguous, while a markedly smaller fraction of unspliced (nascent) reads are classified as ambiguous. This makes sense given that one would expect spliced (mature) molecules to give rise to a substantial fraction of exonic reads. However, this suggests that the proposed assignment rule may still be likely to allocate reads that are truly of spliced (mature) origin as ambiguous, and therefore include them in the context of single-nucleus RNA-seq processing.

We also evaluated how the implied read classifications under each method concord with the proposed model, using the approach described in Section 4.2. While we adopt a different approach to demonstrate the classification agreement (i.e., by showing a multi-class contingency for each method), we find that the results of `kallisto(D-list)|bustools` and `alevin-fry(splici)` in this experiment are largely similar to what is reported by Hjörleifsson and Sullivan *et al*. (11). We find that, by simply replacing the collapsed intronic regions used in `alevin-fry(splici)` with the full-length unspliced (nascent) transcripts in the `alevin-fry(spliceu)` index, we recover mapping results and subsequent classification rules that agree with those proposed by Hjörleifsson and Sullivan *et al*. (11) and underlying this simulation (**Fig. 3)**. Moreover, we observe that the results generated using `alevin-fry(spliceu, piscem)` (i.e., using `piscem` as the underlying mapper) are also highly concordant with those of `alevin-fry(spliceu)`, with only very small difference arising from tweaked mapping heuristics used within `alevin-fry(spliceu, piscem)`. As a result, we see that `kallisto(D-list)|bustools`, `alevin-fry(spliceu)` and `alevin-fry(spliceu, piscem)` demonstrate a high degree of *internal consistency* in their ability to classify reads according to the rules proposed by the assumed model. On the other hand, by adopting a different set of conventions, the mappings from `alevin-fry(splici)` tend to attribute exonic reads (ambiguous under the proposed model) to spliced transcripts, leading to a disagreement with the other methods.

### 2.2. Analysis of experimental single-nucleus RNA-seq data

We compared the quantification results of `kallisto(D–list)|bustools`, `alevin-fry(splici)`, `alevin-fry(spliceu)` (here, we adopted `piscem` as the mapper, though results using salmon, as in section 2.1, are nearly identical), and `STARsolo` on two experimental single-nucleus RNA-seq datasets. As there was a discrepancy between the dataset listed in the manuscript of Hjörleifsson and Sullivan *et al*. (11) ([Adult mouse brain](https://www.10xgenomics.com/resources/datasets/5k-adult-mouse-brain-nuclei-isolated-with-chromium-nuclei-isolation-kit-3-1-standard)) and the files used in the associated scripts ([E18 mouse brain](https://www.10xgenomics.com/resources/datasets/5-k-mouse-e-18-combined-cortex-hippocampus-and-subventricular-zone-nuclei-3-1-standard-6-0-0)), referenced from https://github.com/pachterlab/HSHMP_2022/blob/cfd6958be6ab65fc340bf1df5f1b0e77f19ff967/real_data/README.md?plain=1#L91, we processed both of these datasets. Overall, the relevant results of the analyses comparing different methods are highly-concordant among both of these datasets.

Hjörleifsson and Sullivan *et al*. find that, when processing single-nucleus data, they observe results that are “significantly different from current approaches” (i.e. from `alevin-fry(splici)` and `STARsolo`). Yet, the magnitude of the demonstrated discrepancy appears large, given that these datasets have only a relatively small fraction of unambiguously spliced reads (**Fig. 4)**. In the E18 mouse brain dataset, only about 10% of UMIs are attributed to spliced molecules, and in the adult mouse brain dataset, only about 7% of UMIs are attributed to spliced molecules. Indeed, when we processed these data, we observed *strong agreement* between *all* of the approaches (i.e., both those that adopt the newly-proposed model— `kallisto(D-list)|bustools` and `alevin-fry(spliceu)` — and those that aggregate all counts — `alevin-fry(splici)` and `STARsolo`); see **figures 5 and 6.** Given the distribution of UMI categories, this is not unexpected. The gene-wise UMI correlations are all quite high (Spearman correlation coefficient of 0.88 or above, and Pearson correlation coefficient of 0.98 or above), and the cell-wise UMI correlations are even higher (0.98 or above for both Spearman and Pearson correlation coefficients). This disagrees substantially with the numbers reported by Hjörleifsson and Sullivan *et al*..

**Fig. 5.**
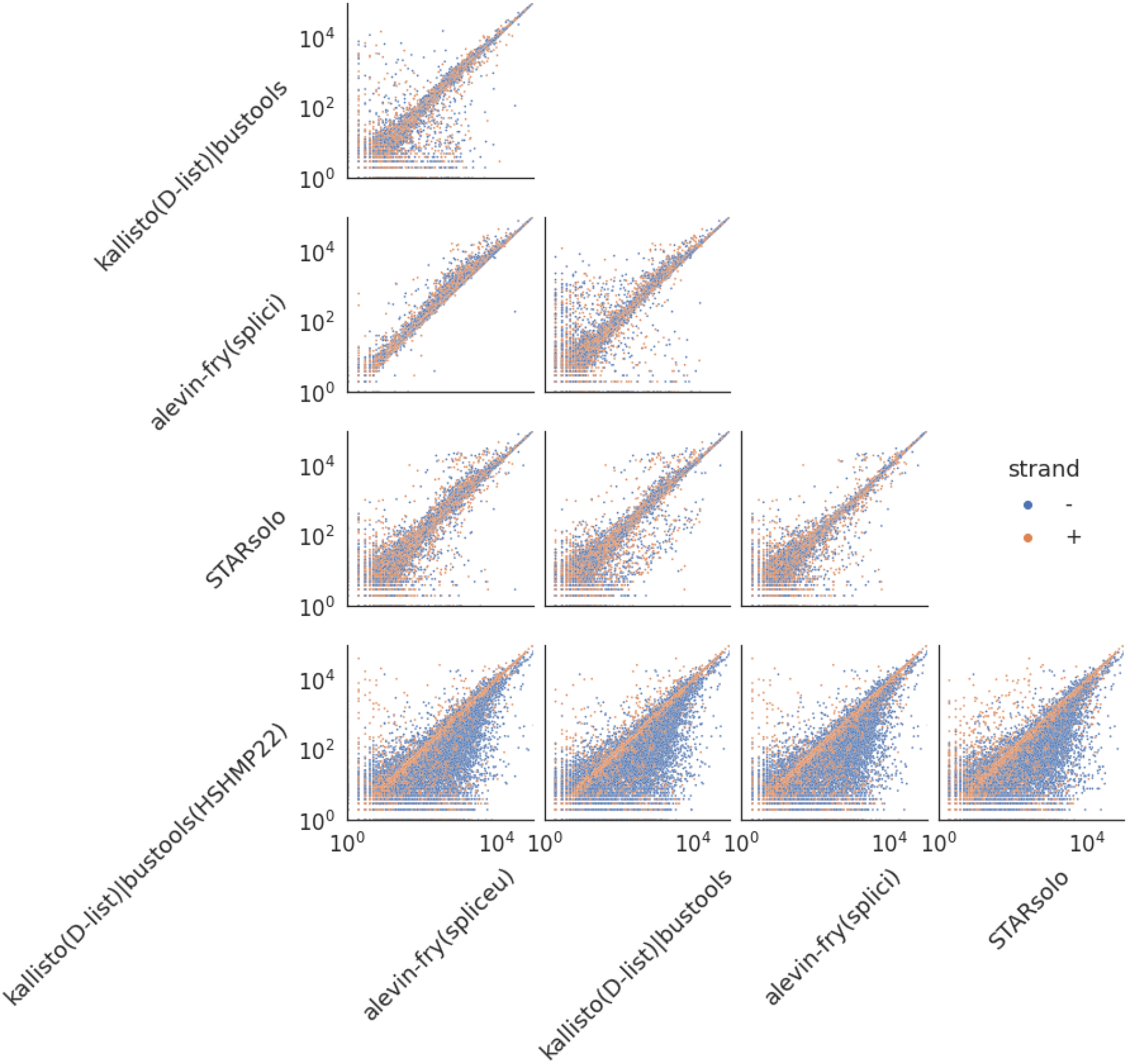
Pairwise scatter plots (on log-scaled axes) of the total UMI count of each gene across all high-confidence cells for each pair of the tested methods on the E18 mouse brain nuclei dataset. The total UMI count of each gene is obtained by taking the sum of the genes’ UMI count across all high-confidence cells from the gene count matrix. In the scatter plots, each dot represents a gene, the value represents the UMI count of each gene. The dots are colored according to the genomic strand on which the corresponding gene resides.

**Fig. 6.**
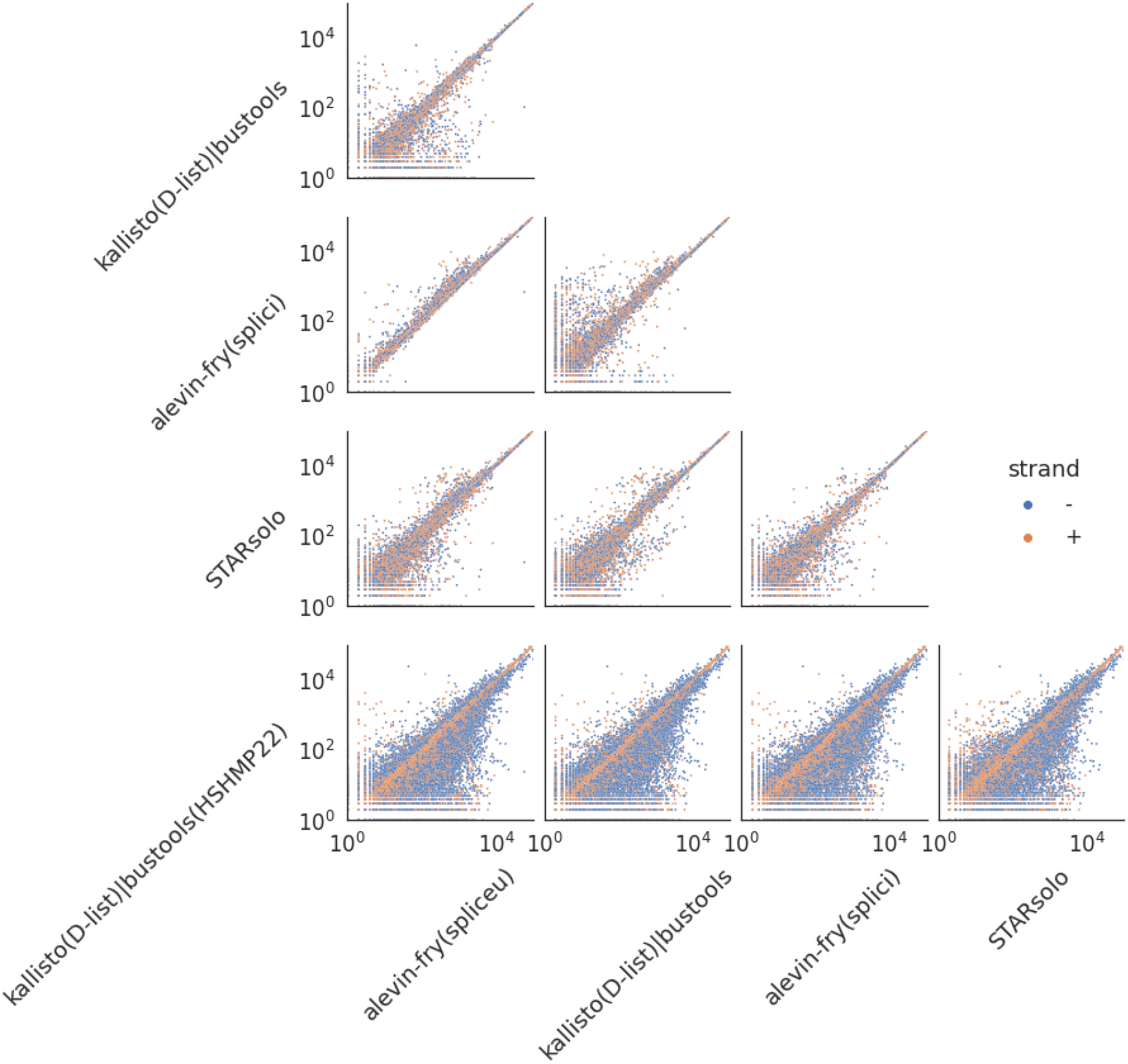
Pairwise scatter plots (on log-scaled axes) of the total UMI count of each gene across all high-confidence cells for each pair of the tested methods on the adult mouse brain nuclei dataset. The total UMI count of each gene is obtained by taking the sum of the genes’ UMI count across all high-confidence cells from the gene count matrix. In the scatter plots, each dot represents a gene, the value represents the UMI count of each gene. The dots are colored according to the genomic strand on which the corresponding gene resides.

To this end, we sought to further investigate what might cause the discrepancy in analysis results. We observed that while the agreement between `alevin-fry(splici)` and `STARsolo` closely matched was reported in (11), the agreement between `kallisto(D-list)|bustools` and other methods did not. One difference in how the data was processed is that we used the same set of unspliced transcripts, extracted as described in section 4, as the reference to build the unspliced (nascent) index for `kallisto(D-list)|bustools`, and as the set of unspliced transcripts in the index for `alevin-fry(spliceu)`. However, to build the unspliced index for `kallisto(D-list)|bustools`, the scripts provided by Hjörleifsson and Sullivan *et al*. (11) reference a file named [mus_musculus_nascent_v2.fa](https://github.com/pachterlab/HSHMP_2022/blob/cfd6958be6ab65fc340bf1df5f1b0e77f19ff967/real_data/README.md?plain=1#L31). Unfortunately, this file is not included in the repository or elsewhere linked, and there is no description of how it was generated. As such, we attempted to use the [generate_cDNA+introns.py](https://github.com/pachterlab/HSHMP_2022/blob/cfd6958be6ab65fc340bf1df5f1b0e77f19ff967/extract_introns/generate_cDNA%2Bintrons.py) script provided in the repository to generate the unspliced transcripts from the mouse genome and annotation file (Ensembl release 108 (18)). This is, for example, how the unspliced transcripts were generated in the simulated classification experiment carried out by the authors. This script produced a substantially different unspliced transcriptome file than what was generated by the procedure we describe in section 4. Specifically, compared to the expected unspliced sequences, the sequences are all one nucleotide shorter, and unspliced sequences arising from genes annotated on the negative strand of the genome are not reverse complemented. This, in turn, results in an unspliced index in which the unspliced transcripts of positive-strand genes are represented mostly correctly (albeit one base too short), while the unspliced transcripts of negative-strand genes are instead represented by their reverse complement. This has a *substantial* effect on the data processing, as the 10x Chromium Next GEM Single Cell 3’ Reagent Kits v3.1 (Dual Index) protocol used in both of these datasets is stranded, and all of the tested tools are executed in configurations that discard reverse-complement strand mappings (i.e., the tools respect the expected strandedness of the protocol).

When processing the data using `kallisto(D-list)|bustools` with this unspliced transcriptome (referred to as `kallisto(D-list)|bustools(HSHMP22))`, we observe results that are, indeed, “significantly different from current approaches.” While these results still do not exactly recapitulate what is reported in (11), they are much closer to what is reported (e.g., the calculated Spearman correlations of gene-level UMI counts are within 0.017 of what is reported, and the difference in Pearson correlations drops from ≥ 0.35 to ~0.1—0.15). While understanding the true cause of the difference between the analyses is not feasible without access to the `mus_musculus_nascent_v2.fa` file, and information about how it was generated, this observation is consistent with the fact that, perhaps, this file was generated using some version of the annotation and the `generate_cDNA+introns.py` script.

Overall, we observe strong agreement between the results of `kallisto(D-list)|bustools` and prior approaches when the indexed nascent transcripts are extracted respecting the annotated orientation. While one may expect divergence to vary between the proposed approach and prior approaches (that include UMIs from molecules of all splicing statuses), based on the fraction of confidently spliced molecules present in the sample being processed, the aggregate differences are quite mild for these data and the methods are largely concordant. In addition to the aggregate statistics of the total per-gene UMI counts, we also evaluated the finer-grained agreement between the approaches by looking at the distribution over all high-confidence cells, of the Spearman correlation of gene-level UMI count vectors (**Figures 7 and 8)**. Here, we observe that all methods evaluated under the same unspliced transcriptome retain relatively high concordance, and that the concordance between the results of those methods and the results of `kallisto(D-list)|bustools(HSHMP22)` is much lower (**Fig. 7)**. Finally, as we expect, by virtue of adopting a *spliceu* reference, we observe that the concordance between between `kallisto(D-list)|bustools` and `alevin-fry(spliceu)` is among the highest between different tools (**Fig. 8)**, since they adopt (implicitly) the same model for read status classification. This demonstrates that, while generally small in these datasets, there is an effect of discarding the counts that arise from UMIs associated with reads that can be confidently classified as spliced (mature).

**Fig. 7.**
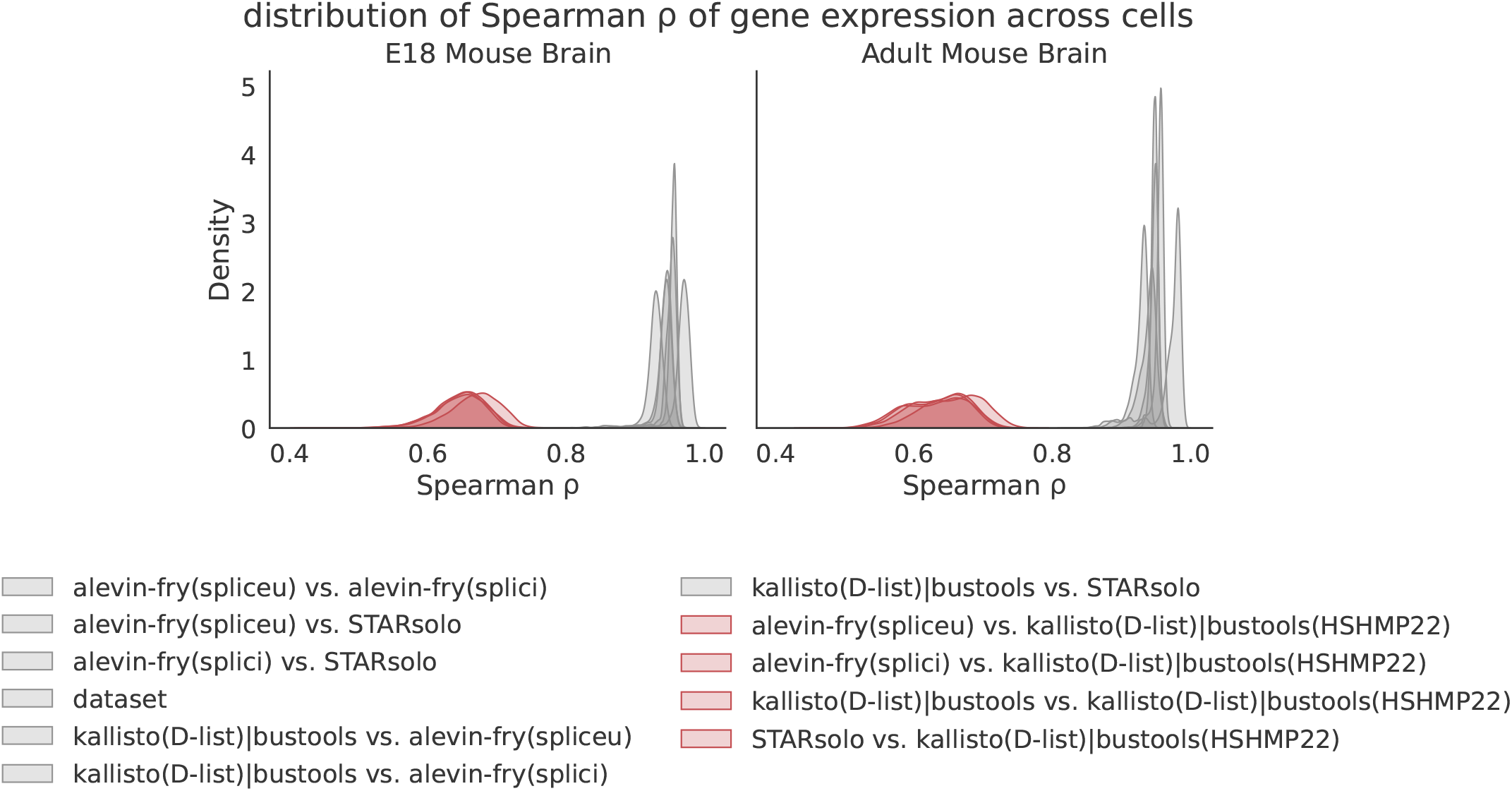
The gene-level Spearman correlation distribution across cells of each pair of the quantification result of the tested methods. The distributions that are colored as red represent the comparison of all methods with the `kallisto(D-list)|bustools` result using the nascent transcripts extracted by following the `generate_cDNA+introns.py` script from Hjörleifsson and Sullivan *et al*. (11). The left plot is for the E18 mouse brain nuclei dataset, and right plot is for the adult mouse brain nuclei dataset.

**Fig. 8.**
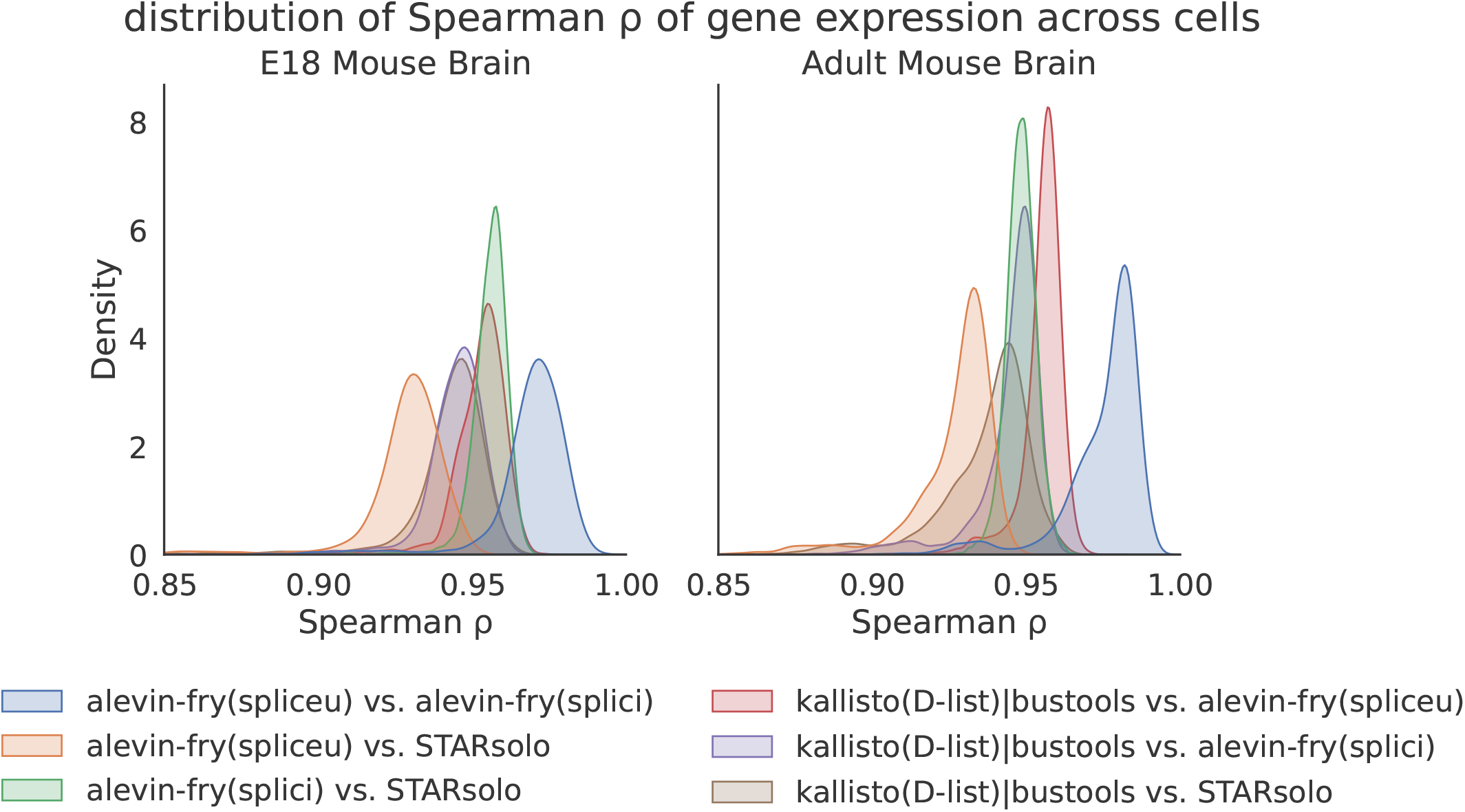
The cell-level Spearman correlation distribution across genes of each pair of the quantification result of the tested methods, sans `kallisto(D-list)|bustools(HSHMP22)` (which is an outlier).

### 2.3. Possible paths forward: Incorporating probabilistic evidence into the inference of UMI splicing status

While complete inference of the splicing status of the molecules that generate observed reads is a very challenging task, the discrete classification of each UMI as unspliced, spliced, or ambiguous is only as useful as the ability of downstream tools to make use of these categories. Ultimately, it is likely the case that, to make optimal use of this data, either (i) downstream tools will have to incorporate UMIs of ambiguous splicing status into their models and processing, or (ii) quantification tools will have to adopt more comprehensive inference to provide evidenced-based predictions of splicing ambiguity (or both). When downstream methods do not explicitly model splicing-ambiguous UMIs, leaving UMIs classified as ambiguous, allocating all ambiguous UMIs to one or another of the definitive splicing statuses, or even adopting any fixed fractional allocation, are all suboptimal solutions.

We believe that it will be useful to build a probabilistic model for the splicing status (and gene-level) ambiguity of sequenced molecules in tagged-end single-cell RNA-sequencing, as has been done successfully for the problem of transcript-level abundance estimation in bulk RNA-seq analysis (17, 19–22). While developing a complete probabilistic generative model and implementing inference over such a model is a substantial and non-trivial challenge that has not yet been resolved, there are nonetheless efforts underway to build models of splicing dynamics (23) that would allow estimation of spliced and unspliced molecule ratios, though such models can be complex and may not be able to be robustly estimated for all genes or across all cellular contexts. Likewise, models of UMI generation are also being developed that allow probabilistic resolution of multi-gene associated UMIs (24), and previous work has demonstrated that an empirical Bayesian approach can even be used to share information among cells within a sample to help resolve UMI assignment (25). Possibly, such improved models could even be built atop existing methods that already seek to perform probabilistic UMI assignment, like the `parsimony-em` resolution strategy of `alevin–fry` (13,26), if one considers the spliced and unspliced transcripts of a gene as coming from different genes, as was done by Soneson et al. (14) in the context of estimating counts upstream of RNA-velocity analysis. Nonetheless, to improve upon existing models and enable more accurate inference of splicing status and gene-level ambiguity, it will be essential to account for extra information that can be gleaned from the data.

As mentioned above, and in the conclusion, mapping and quantifying against a reference that contains the full set of spliced and unspliced transcripts can prove useful in recovering some ambiguous UMIs. For example, if a UMI is associated with multiple reads where some are of ambiguous splicing status and others are of definitive splicing status, the reads of definitive status can be used to “rescue” the ambiguous reads and assign the UMI a definitive status. Nonetheless, due to the prevalence of purely exonic reads, many UMIs are likely to remain ambiguous under conservative assignment models. Here, incorporating other evidence to help assign a probabilistic splicing status may be useful.

In the evaluation of the splicing status of UMIs, one piece of potentially useful evidence that is currently unused is the likelihood that particular priming sites give rise to the reads associated with a UMI. For example, since the priming of sequenced fragments is expected to originate from the priming of poly(dT) primers to adenine single-nucleotide repeats (A-SNRs) (27) along transcripts, and since fragment lengths in tagged-end single-cell RNA-seq follow a well-characterized distribution that can be inferred from the observed data, one can evaluate the likelihood that a particular A-SNR gives rise to an observed fragment based on the mapping location of the “biological” read. Specifically, for a read with a given mapping position x on a transcript t, one can evaluate the probability that the associated fragment pairs this mapping with every A-SNR downstream of x by evaluating the probability of observing a fragment of the implied length under the empirical fragment length distribution in this sample. If it is highly likely that a read is paired with an A-SNR located within an intron, then this provides strong evidence that the associated UMI should be assigned an unspliced status. On the other hand, if the read is likely paired with the terminal A-SNR (i.e., the poly-A tail), then the pairing is not particularly informative as to the associated UMI’s splicing status.

More formally, let the fragment length distribution of the current sample be *f*. This is a probability distribution that assigns a probability *p_f_*(*l*) to observing a fragment of length *l*. Further, for a read mapping to position *x* of exon *e* on gene *g*, let asnr(*g,e,x,s*) = {(*a_1_*,*t_1_*,*d_1_*),…,(*a_j_*,*t_j_*,*d_j_*)} denote the set of *all* A-SNRs downstream of this position in *any* spliced transcript of *g* that contains e. Here, ar is an A-SNR site, and di is the associated distance from the fragment start position to this site along transcript ti. Likewise, let asnr(*g,e,x,u*) be analogously defined as the set of *all* A-SNRs downstream of position *x* on exon *e* of gene *g* in any unspliced transcript of *g* that contains *e*. Given a read *r* of ambiguous status that maps to position *x* on exon *e* of gene *g*, we can evaluate the probability that this read arises from a spliced transcript as:

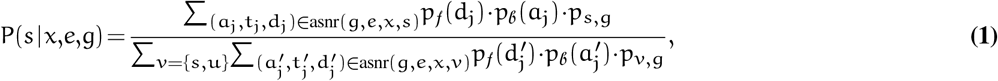

where *p_b_*(*a_j_*) is a “binding” model that assigns a binding probability to an A-SNR (or near A-SNR) motif, since sufficiently long stretches of adenine, even with mismatches, may still bind to poly(dT) primers associated with the sequenced fragments (28). The terms *p_s,g_* and *p_u,g_* are prior expectations of observing the spliced (respectively unspliced) transcripts of gene *g*, and could be set to be uninformative, or could be informed and iteratively estimated within the context of a more comprehensive model. While this only provides a way to evaluate the splicing status probability for an individual read and not a UMI, one could adopt a simple approach to aggregate this read-level information to the UMI level by taking the UMI-associated probability to be the expectation of the probabilities of the associated reads — that is, for a UMI with an associated set of m read mappings 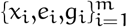:

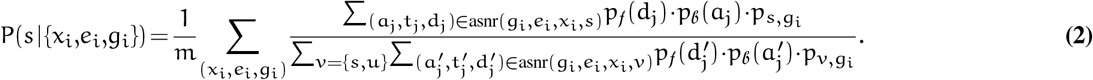

If the read mappings associated with this UMI are all to the same gene *g*, then it will be the case that (*x_i_*,*e_i_*,*g_i_*) = (*x_i_*,*e_i_*,*g*). If the UMI has read mappings associated with multiple genes, then one may discard such a UMI, first assign it to one of the mapped genes based on another set of rules or heuristics, or incorporate the above splicing probabilities into a multi-gene UMI resolution model, where each mapping can be weighted by the estimated (and iteratively updated) expectation that the UMI is associated with each of the multi-mapped genes; this simply changes the manner in which the expectation is calculated.

While the development of a complete probabilistic model for splicing (and multi-gene) UMI resolution is a challenging task, it is likely to be a fruitful one, and there are several pieces of evidence that can be used to improve upon the current all-or-nothing or fixed fractional allocation schemes for assigning splicing statuses to UMIs. Here, we have suggested how one particularly informative piece of evidence — the implied fragment length — can be used to help probabilistically resolve UMI splicing status. Though the complete specification and implementation of inference over this model is beyond the scope of the current work, we plan to further develop these ideas and incorporate them into the `alevin-fry` single-cell data processing pipeline.

## 3. Conclusion

Single-cell and single-nucleus RNA-seq data processing has attracted increasing attention in recent years. One challenge to accurate quantification is how to handle ambiguity, which arises in terms of UMIs that are associated with reads that align to multiple genes, as well as reads that are compatible with both the spliced and unspliced transcripts of a gene.

Several heuristic conventions have been proposed to date to assign a splicing status to UMIs for various types of subsequent analysis. Further, the conventions related to how UMIs arising from different splicing statuses should be handled when quantifying different assays are actively shifting as we learn more, as a community, about the nature of the data and the type of information contained within it. For example, recent work suggests potential benefits to including UMIs derived from unspliced molecules or other sources, even when quantifying what is expected to be spliced RNA (1–3). However, even these approaches are unlikely to be the final word about best practices for quantification. We anticipate that further studies and analyses will lead to improved models and methods for accurate and comprehensive quantification, which, in turn, will improve downstream analyses.

At this point, then, it is desirable for preprocessing tools to enable fast and accurate quantification of the UMIs arising from each gene and their associated splicing status, acknowledging that an ambiguous category may be necessary, since a definitive splicing status cannot always be derived from the observed data. Therefore, we contend that retaining this ambiguous category and propagating it to subsequent stages of analysis may be preferable to collapsing the ambiguous category into a definitively spliced or unspliced status, even if that collapse is performed differently for single-cell and single-nucleus RNA-seq data.

Likewise, mapping and quantifying against a reference that represents both the spliced and unspliced transcripts seems preferable to mapping against separate references depending upon the type of data being processed, both because it provides more information to UMI resolution methods, so that some ambiguous UMIs may be properly assigned a definitive status, and because it propagates gene counts for all splicing statuses to downstream analyses. Subsequently, downstream tools can discard the UMIs arising from the “unexpected” splicing status if desired, or they can incorporate this information into their models when it may prove helpful. Moreover, these categorized counts can be obtained with a single mapping and quantification run for improved efficiency and reduced complexity. While existing methods can already index the joint set of spliced and unspliced transcripts in quite moderate memory (13), we have introduced a further-improved index and mapping tool, `piscem` (16), that can be used directly upstream of `alevin-fry`, and that is capable of mapping against the joint spliced and unspliced transcriptome in ≤ 3GB of memory for human and mouse references. We also believe that a fruitful direction for future research will be developing a comprehensive generative model for sequenced reads and UMIs that makes the best use of available information (including e.g., implied fragment lengths) to resolve both gene-level and splicing-level ambiguity of UMIs probabilistically.

In our study, we acknowledge that there are limitations and relevant questions left unexplored or underdeveloped. For example, when constructing the *spliceu* transcriptome, we have only evaluated the inclusion of a single nascent transcript per gene, rather than considering all distinct unspliced precursors of annotated spliced transcripts (14). The latter approach will induce more multimapping ambiguity, but can also represent more sequences and allow more accurate determination of the position of mapped reads on the transcripts from which they might originate. Further, we have proposed how additional evidence can be used to provide improved predictions of splicing ambiguity, but have not yet fully incorporated these ideas into a formal probabilistic generative model or implemented them in practice. Finally, we have focused, here, entirely on tagged-end single-cell technologies. However, alternative technologies (e.g. (29, 30)) may avoid strong positional biases and provide substantially different evidence regarding splicing status to preprocessing tools, and may therefore warrant their own models and methods. These are all interesting directions for future work.

## 4. Methods

### 4.1. Extracting mature and nascent transcripts

To build `alevin-fry` <u>spliced</u> and <u>u</u>nspliced (*spliceu*) indices and `kallisto(D–list)|bustools` indices with D-lists, it is necessary to extract the spliced (mature) and unspliced (nascent) transcripts from the genome and gene annotation. We observed that the Python script provided by Hjörleifsson and Sullivan *et al*. (11), [generate_cDNA+introns.py](https://github.com/pachterlab/HSHMP_2022/blob/89e2e3b68eb6747f28aac1728a8268f39c9eba47/extract_introns/generate_cDNA%2Bintrons.py), does not properly extract the annotated transcript sequences, as it neither considers the strandedness of the unspliced transcripts, nor extracts the complete sequence of genomic units. More precisely, when extracting a spliced transcript by concatenating the exons of the transcript, it extracts all but the last base of each exon from the forward strand of the corresponding chromosome; when extracting the unspliced transcript of a gene, it extracts all but the last base of the gene body with respect to the forward strand of the chromosome. Further, the sequences representing the spliced and unspliced transcripts are written as extracted, regardless of their strand of origin. The effect of this issue appears to be non-uniform in the analyses carried out by Hjörleifsson and Sullivan *et al*. (11). For example, it affects the transcripts extracted for use in the classification experiment on simulated data, but the magnitude of that effect is small since all tools are run in a strand-unaware fashion, largely offsetting the fact that transcripts from negative strand genes are not reverse-complemented. On the other hand, the effect on the analysis of experimental single-nucleus data may be substantial (see Section 2.2).

In this work, spliced transcripts were made by concatenating the sequence of exons according to the annotated strand using the `extractTranscriptSeqs` function from the `GenomicFeatures` (31) Bioconductor package. The `genes` function from `GenomicFeatures` and `getSeq` function from `BSgenome` (32) were used to define and extract a single nascent transcript per gene. Finally, extracted spliced and unspliced transcripts were combined to make the *spliceu* reference. During this process, the gene annotation GTF file was loaded as a TxDb object using the `makeTxDbFromGFF` function from `GenomicFeatures`, and the genome FASTA file was loaded as a `DNAStringSet` object from the `Biostrings` (33) Bioconductor package. Note that the “unspliced transcript” of a gene defined by Hjörleifsson and Sullivan *et al*. (11), which is also applied in the *spliceu* reference construction in this work, actually represents the whole gene body, instead of the unspliced precursor of each spliced mRNA, i.e., the genomic range from the start of the first exon to the termination of the last exon (14). The drawback of defining one unspliced transcript per gene is that, for reads originating from unspliced transcripts, one immediately loses the position information of the mapping site with respect to the unspliced transcript origin, which might be helpful for resolving the ambiguity (as suggested in Section 2.3). To avoid this, one can easily construct a *spliceu* reference that contains one unspliced transcript for each spliced transcript, as defined in Soneson et al. (14), though that is not the convention we follow in this work.

### 4.2. Classification Simulation

Hjörleifsson and Sullivan *et al*. (11) introduced a new simulation focused on assessing the concordance between existing read assignment strategies and the classification scheme they propose. This simulation generated reads from nascent and mature transcripts to mimic the distributions of nascent and mature molecules sequenced in both the cytoplasmic and nuclear contexts. The simulated reads in both contexts are then converted into a 10x Chromium v3 compatible FASTQ format, with randomly assigned barcodes (a 10x barcode and a unique molecule identifier). The mapping result returned by each method is then individually inspected to assess the assignment of each read.

This simulation experiment uses `bbmap` to simulate reads without error. We note that it also elects to leave certain, potentially relevant properties of tagged-end scRNA-seq data unmodeled. For example, the proximity of generated reads to primable sites is not accounted for in the simulation, so the positional origin of reads with respect to the transcripts from which they are generated is unlikely to mimic what would be observed in experimental data. Likewise, while the 10x Chromium protocol is strand-aware, the simulated data are generated in an unstranded fashion, and therefore produce substantially more anti-sense reads than one would expect to observe in experimental data (1). This lack of strand-awareness is also likely to artificially inflate the mapping ambiguity of the simulated reads.

Finally, while certain scripts related to the original classification experiment were available in the repository accompanying the manuscript of Hjörleifsson and Sullivan *et al*. (11), several referenced files and scripts were missing, making precise replication of that simulation experiment impossible. Therefore, we instead chose to reimplement this simulation following the description in the manuscript and the script files that were available as closely as possible, except where otherwise noted.

#### Sequencing read simulation

The spliced and unspliced transcripts in the *spliceu* reference, extracted from the [(GRCh38) version 2020-A](https://support.10xgenomics.com/single-cell-gene-expression/software/downloads/latest) human reference, were used to invoke the `randomreads` program of `bbmap` to simulate cytoplasmic and nuclear datasets. Both datasets have five million reads of length 150. The biological reads in the cytoplasmic dataset were generated by simulating four million reads from the mature transcripts from both strands, and one million reads from the nascent transcripts from both strands. Similarly, the biological reads in the nuclear dataset were generated by simulating one million reads from the mature transcripts, and four million reads from the nascent transcripts.

Each simulated biological read was assigned a corresponding technical read, which consists of a random 10x barcode of length 16 and a random unique molecule identifier (UMI) of length 12. The simulated read pairs were then packed into the FASTQ format, so that they can be processed by the tested methods as a 10x Chromium v3 *3’* dataset.

#### Determination of ground truth read class

In the simulated data, every generated read has a definitive splicing status according to its true origin (i.e., the transcript from which it was truly simulated). Thus, there are no ambiguous reads with respect to the simulated locus of origin. Rather, to assess the ability of different approaches to classify each read based on the collection of mature and nascent transcripts that match the read, and the classification rules being evaluated, this experiment assigned each read one of three possible classes: mature, nascent, or ambiguous. Since all reads were simulated without error, all matches are perfect and equally optimal, and thus reasonable mapping disambiguation cannot be performed for each read in isolation (i.e., the status of other reads sharing the same UMI and barcode can not be used to disambiguate a read). In this sense, the current experiment assesses the *internal consistency* of the mapping result with the mature, nascent, and ambiguous classification scheme proposed by Hjörleifsson and Sullivan *et al*. (11). To generate a *ground truth* class for each simulated read in a manner consistent with what they proposed, we used the following procedure.

A (multi-string) suffix array index (based upon (34)) was constructed over all unspliced and spliced transcript sequences. Each simulated read (and its reverse complement) was matched against this suffix array to determine all locations where the read was a substring of the reference. If the read matched only mature transcripts, then it did not come from a region of mature transcripts that was shared with any nascent transcript, and it was assigned a spliced (mature) status. If the read matched only to unspliced transcripts, then it did not come from a region of unspliced transcripts that was shared with any spliced transcript, and it was assigned an unspliced (nascent) status. Finally, if the read matched both spliced and unspliced transcripts (possibly of different genes), then it was assigned an ambiguous status, since the mapping information alone is inadequate to assign this read a definitive splicing status.

Again, it bears repeating that this experiment focuses on evaluating the *internal consistency* of the mapping results produced by each method with the classification scheme proposed by Hjörleifsson and Sullivan *et al*. (11). Therefore, what it is primarily evaluating is the ability of each method’s combination of index and mapping algorithm to produce mapping results that accord with the proposed assignment rule.

#### Read alignment

The simulated reads were aligned to the human reference (GRCh38) version 2020-A, the same reference used for generating the simulated reads, using `STARsolo`, `kallisto(D-list)|bustools`, and `alevin-fry` with both the *splici* and *spliceu* references. For all variants of `alevin-fry`, mapping was performed using pseudoalignment (22) with structural constraints (13) corresponding to the `--sketch` flag in `salmon alevin` (likewise, this is the mapping algorithm implemented in `piscem`). The `STARsolo` index was constructed using the matched human genome reference and annotation. The `alevin-fry` indices were built on the *splici* and *spliceu* references made from the human reference. The `kallisto(D-list)|bustools` index was constructed on the mature transcripts made by calling the `kb ref` command from `kb-python` (35) on the human reference, with the whole genome as the D-list. The nascent index was constructed on the same nascent transcript set used to build the `alevin-fry(spliceu)` reference, and the spliced transcriptome was provided as the D-list. After constructing the indices for the tools, read mapping of each tested method was performed by following the commands in the scripts provided by Hjörleifsson and Sullivan *et al*. (11).

#### Determination of empirical read class

The procedure to assign an empirical read class differs for genome-based and transcriptome-based methods. As each read has a unique barcode (10x barcode plus UMI), barcode sequences reported in the mapping results were used to identify the simulated reads.

For transcriptome-based tools, including `kallisto(D-list)|bustools` and `alevin-fry`, the empirical class of each simulated read is assigned according to the compatible reference transcripts in the corresponding mapping result. Using the same strategy as in Section 4.2, the read was assigned a mature status if it matches only mature transcripts, nascent status if it matches only nascent transcripts, and ambiguous otherwise. As was done by Hjörleifsson and Sullivan *et al*. (11), the full quantification algorithm was not run. Rather, the intermediate mapping results were programmatically evaluated to infer the implied status. As the reads were simulated from both strands of the underlying reference sequences, the mappings were not filtered by strand before being used to determine the empirical classification.

Unfortunately, since `STARsolo` aligns the reads in a (potentially) spliced fashion to the genome, the same simple strategy cannot be used to easily assign an inferred status to each read from the results produced by `STARsolo`. It also seems there is not a simple way to obtain this read-level information from the intermediate files produced by `STARsolo` (personal communications with Dr. Alex Dobin). Thus, rather than following a more complex procedure, we elected to elide the evaluation of `STARsolo` in this particular experiment.

#### Evaluation metrics

To compare the empirical read class inferred for each method with the matching-based ground truth, we computed a multi-class contingency table. This demonstrates, in each context (cytoplasm and nuclear), the frequency with which reads having a certain matching-based ground truth class under the classification rules being evaluated are assigned to a different class. Moreover, in this simulation, as the splicing status of the transcript of origin of each read is given by `bbmap`, we also compared the splicing status of the transcript origin of each read with the matching-based ground truth of the read.

To provide a concise yet comprehensive summary of agreement between the model and the different methods (as well as between the model and the true splicing origin), we chose to compare the the splicing status of each method against the matching-based ground truth using a multi-class contingency table. Likewise, we use the same approach to compare the matching-based ground truth against the true origin, which lacks an ambiguous category. Each table is represented as a heatmap in **Figures 2 and 3.** These heatmaps show how frequently a read deemed to belong to a particular category based on the mappings of a method agrees with the matching-based ground truth classification (**Fig. 3)**, and how frequently the matching-based ground truth is able to recover the true splicing origin versus declaring a read as ambiguous (**Fig. 2)**.

### 4.3. Analysis of mouse single-nucleus RNA-seq data

In this experiment, we analyzed two different single-nucleus RNA-seq datasets. First, we analyzed a [5k Mouse E18 Combined Cortex](https://www.10xgenomics.com/resources/datasets/5-k-mouse-e-18-combined-cortex-hippocampus-and-subventricular-zone-nuclei-3-1-standard-6-0-0), [Hippocampus and Subventricular Zone Nuclei](https://www.10xgenomics.com/resources/datasets/5-k-mouse-e-18-combined-cortex-hippocampus-and-subventricular-zone-nuclei-3-1-standard-6-0-0) dataset from 10x Genomics using the mouse GRCm39 (Ensembl 108) reference downloaded from [Ensembl](http://asia.ensembl.org/Mus_musculus/Info/Index) (18). According to the dataset’s web page, there are 5,973 detected cells. This is different from the number reported in Hjörleifsson and Sullivan *et al*. (11), but this dataset matches the file names of the FASTQ files in the [associated commands](https://github.com/pachterlab/HSHMP_2022/blob/cfd6958be6ab65fc340bf1df5f1b0e77f19ff967/real_data/README.md?plain=1) in the GitHub repository. We also analyzed the [5k Adult Mouse Brain Nuclei Isolated with Chromium Nuclei Isolation Kit](https://www.10xgenomics.com/resources/datasets/5k-adult-mouse-brain-nuclei-isolated-with-chromium-nuclei-isolation-kit-3-1-standard) dataset from 10x Genomics, which is the dataset mentioned by Hjörleifsson and Sullivan *et al*. (11), containing 7,377 detected high-confidence cells.

The sequencing reads were aligned to the mouse GRCm39 108 reference using `STARsolo`, `kallisto(D-list)|bustools`, and `alevin-fry` with the *splici* and *spliceu* references. For all variants of `alevin-fry`, including downstream of `piscem`, mapping was performed using pseudoalignment (22) with structural constraints (13). A STARsolo index was constructed using the mouse reference. Al evin-f ry indices were constructed using the *splici* and *spliceu* references extracted from the mouse reference. For counting UMIs in these experiments using `alevin-fry` in accordance with the rules proposed by Hjörleifsson and Sullivan *et al*., after quantifying with the *spliceu* index, the unspliced and ambiguous layers of the resulting USA-mode count matrix (13) were summed to produce the main count layer used for comparison. A `kallisto(D-list)|bustools` index was constructed using the unspliced (nascent) transcripts in the `alevin-fry` *spliceu* reference, with the mature transcripts generated by calling the `kb ref` command on the mouse reference as the D-list. Additionally, in a further attempt to reproduce the results reported in Hjörleifsson and Sullivan *et al*. (11), another `kallisto(D-list)|bustools` index was constructed using the mature transcripts obtained using `kb ref`, and the (strand-unaware) unspliced (nascent) transcripts extracted by the `generate_cDNA+introns.py` script. Using these indices, we followed the steps in Hjörleifsson and Sullivan *et al*. (11) to quantify the dataset. The relevant metrics over cells and genes were computed and reported using the quantification results of the tested methods.

### 4.4. The piscem index

`Piscem` (16) is an improved index for the compacted reference de Bruijn graph that admits the same set of operations as the `pufferfish` (15) index, but that is substantially more space efficient. Abstractly, the index is characterized as the composition of a k-mer-to-tile map and a tile-to-occurrence map. By composing these two maps, one obtains a fully-functional k-mer-to-occurrence index. `Piscem` uses `cuttlefish` (36) to efficiently construct the compacted reference de Bruijn graph and associated tiling. It then uses `sshash` (37) as an efficient k-mer-to-tile map, and it builds a dense (but compacted) inverted index to represent the tile-to-occurrence map. This tile-to-occurrence map can be further compressed using the methodology developed by Fan et al. (38), though this feature has not yet been implemented in the software. A complete manuscript describing the `piscem` index, mapping algorithm, and software implementation, and its application beyond the mapping of single-cell RNA-seq data, is currently in preparation.

`Piscem` can be used as a drop-in replacement for `salmon` (17) to perform mapping upstream of `alevin-fry` (13). The `piscem` index is *considerably* smaller than either the dense or sparse `pufferfish` indices. For example, the *full spliceu* indices, containing all nascent and spliced transcripts, are 1.8GB and 1.9GB, respectively, for human (GRCh38-2020-A) and mouse (GRCm39.108). Further, the structure of the index on disk is essentially the same as in memory. This allows the `piscem` single-cell mapper to operate in a very low memory footprint; ≤ 3GB for all of the data evaluated in this manuscript. Therefore, the `piscem` index and mapper represent an enticing option when memory is at a premium. Likewise, this is the index around which future development will be performed, as it allows scaling to much larger reference sequences (e.g., full genomes, large genome, and metagenome collections) while requiring only moderate memory resources.

#### Associated scripts

The scripts used to produce the results presented in this manuscript are roughly based on those accompanying Hjörleifsson and Sullivan *et al*. (11) and are available at https://github.com/COMBINE-lab/scrna-ambiguity.

#### Software information

In the analyses performed in this work, we used salmon 1.9.0, `alevin-fry` 0.8.0, `STAR` 2.7.10b, `kb_python` 0.27.3, `piscem` 0.3.0, `BBMap` 39.01, `simpleaf` 0.8.1, `empirical_splice_status` 0.1.0, `kallisto-D` 0.48.0 (commit Cfd6958be6ab65fc340bf1df5f1b0e77f19ff967), `bustools` 0.42.0 (built from commit `89a8a5ddee4ec1c3bb95dfb42e13f2d2ba86e9a1`), R 4.1.3, `BSgenome` 1.62.0, `GenomicRanges` 1.46.1, `doParal-lel` 1.0.17, and `Biostrings` 2.62.0.

## Acknowledgments

The authors would like to thank Mike Love and Stephanie Hicks for useful feedback and discussions related to this work and manuscript.

